# Simultaneous production of lipopeptide and rhamnolipid biosurfactants by *Pseudomonas aeruginosa:* A promising blend for biosurfactant-enhanced bioremediation

**DOI:** 10.1101/2024.10.15.618535

**Authors:** María del Rosario Jacoby, Marcos E. Dening, Laura J. Raiger Iustman

**Author notes:** <u>**Corresponding author address:**</u> Laura J. Raiger Iustman: Ciudad Universitaria, Pabellón 2 4th floor, laboratory QB47. (z.c.1428). Ciudad Autónoma de Buenos Aires. Argentina. Maria del Rosario Jacoby present address: Department of Ecology, Genetics and Evolution, IEGEBA - CONICET, Faculty of Exact and Natural Sciences. University of Buenos Aires.

## Abstract

Oil contamination is a significant environmental issue worldwide, and in the last decades, bioremediation has emerged as a preferred strategy to address this challenge. However, the hydrophobic nature of oil and its limited bioavailability affect its biodegradation by the soil microbiota. To mitigate these limitations, surfactant use has been proposed to enhance oil bioremediation. Nowadays, there is a growing interest in reducing the carbon footprint, and the use of biosurfactants instead of synthetic ones contributes to this global objective. In this study, surfactant-producing bacteria were isolated from a polluted urban stream in Buenos Aires Province to obtain cell-free biosurfactant extracts for use as additives in surfactant-enhanced remediation (SER) protocols in diesel-contaminated microcosms. Out of five isolates, the surfactant extracts of two *Pseudomonas aeruginosa* strains were selected. One of them showed a significant improvement in diesel degradation compared with the controls. Remarkably, this extract was composed of a blend of rhamnolipids and lipopeptides. This is the first work documenting the co-production of both kinds of surfactants by a *P. aeruginosa* strain and its potential for application in surfactant-enhanced bioremediation strategies.

## 1. Introduction

Environmental contamination with petroleum and its derivatives is an ecological threat due to its impact on human and ecosystem health [1]. Hydrocarbons are introduced into the environment through both natural and anthropogenic sources. Depending on the origin of the pollution, hydrocarbon-contaminated environments may contain light petroleum products such as gasoline, kerosene, and diesel; heavy hydrocarbons like lubricants, heavy oil, and crude oil; or complex molecules, including aromatic and polyaromatic hydrocarbons [2]. Diesel is a distillation product of crude oil composed of various hydrocarbons, including straight and branched alkanes, cycloalkanes, and aromatic compounds [3]. The presence of these compounds in soil alters its physicochemical and biological parameters, such as pH, oxygen, and nutrient availability, water-holding capacity, and hydrophobicity. These changes influenced the microbial community structure, including shifts in diversity, activity, and composition [4].To mitigate the impact of these contaminants, different remediation techniques like incineration, landfill, leaching, chemical oxidation, and bioremediation have been proposed [5]. One of the bioremediation strategies, named biostimulation, implies the addition of nutrients to stimulate the native microbiota to enhance pollutant degradation. But despite the biodegradability of hydrocarbons, their limited bioavailability represents a challenge to oil microbial degraders. To address this, the use of surfactant in biostimulation protocols was proposed, leading to the development of Surfactant Enhanced Remediation (SER) [6-7]. Chemical surfactants, produced by industrial synthesis, have been shown to be effective in SER protocols, but also produce negative effects on the environment, including the alteration of soil microbial community and soil enzymatic activity [8]. On the other hand, microbial surfactants or biosurfactants (BS) are less toxic, biodegradable, chemically versatile, and stable at a broad range of pH and temperature [9]. Biosurfactants are amphipathic molecules consisting of a hydrophobic moiety, like fatty acid or its derivatives, and a hydrophilic moiety that could be carboxylic acids, amino acids, carbohydrates, alcohols, or peptides [10].

Among microbial surfactants, lipopeptides and glycolipids are the most commonly studied [11]. Bacterial lipopeptide surfactants are synthesized by nonribosomal peptide synthetases (NRPSs) - a multidomain enzyme that synthesizes oligopeptides without the ribosomal system - leading to the formation of a wide variety of lipopeptides [12]. Those linear or cyclic oligopeptides are then acylated with different fatty acid moieties. These compounds are commonly secreted by bacterial genera such as *Bacillus (*surfactins, fengycins, and iturins), *Streptomyces* (daptomycin), and *Pseudomonas syringae* and *fluorescens* complex (viscosin, amphisin, tolaasin, and syringomycin) [11-12]. Lipopeptides are known for their strong pseudo-solubilization properties and have found applications in environmental protection, industry, and medicine, including antimicrobial and antitumor agents [8]. Among the glycolipid surfactants, the most studied group is rhamnolipids, particularly those produced by *Pseudomonas aeruginosa*. The biosynthesis of rhamnolipids involves three key enzymes: RhlA and RhlB, involved in mono-rhamnolipids synthesis, and RhlC, which adds a second rhamnose to produce di-rhamnolipids. The *rhlA* and *rhlB* genes are located on a single operon regulated by *rhlI* and *rhlR* genes. The third gene, *rhlC*, is also controlled by the *rhlI/R* system but is located in a different locus on the *P. aeruginosa* genome [13].

In SER assays, biosurfactants have been shown to be more effective than synthetic ones, enhancing oil-in-water pseudo-solubilization while maintaining the soil’s biological functions and diversity [14]. Nevertheless, BS production costs make them less competitive than synthetic surfactants, especially because of the costs of the used carbon sources like glucose and glycerol, and the purification process after fermentation [15].

Surfactant post-fermentation production implies several purification steps, including cell-free supernatant acidification, crude product extraction, concentration, and several purification steps like ion exchange, adsorption-desorption, and preparative chromatography [16]. To avoid extra production costs, partially purified biosurfactants were used in several SER assays [15]. Given this approach and the increasing demand for new biosurfactants [17], the objective of this work was to isolate and characterize biosurfactant-producing bacterial strains, to obtain biosurfactant crude extracts (BCEs), and assess their effectiveness in Surfactant-Enhanced Remediation (SER) in soil microcosms.

## 2. Materials and methods

### 2.1. Bacterial strain isolation and cultivation

Water samples for the isolation of diesel-degrading, biosurfactant-producing bacteria were obtained from an urban stream located in the neighborhood of an industrial waste treatment plant in Moreno district, Buenos Aires Province, Argentina (34º 34’ 50.9’’ S, 58º 49’ 25.6’’ W) in December 2017. Sub superficial water was collected with a 5 liters bulk and transferred to an sterile bottle for carrying it to the laboratory and keeping it at 10°C. for the next 48 h. Eight subsamples of 10 ml each were supplemented with 10% v/v filtered (0.22 ◻m-Microclar, Argentina) sterile diesel, 1% mM MgSO_4_.7H_2_O, and 1% MT (micronutrients) solution [18]. When necessary, 0.2 or 0.02 % of yeast extract was added as a source of micronutrients and vitamins. Samples were incubated in 100 ml capped bottles at 30°C and 250 rpm for two to seven days until turbidity development, then 100 µl of each culture were inoculated in 10 ml of E2 minimum media [13] supplemented with 10% v/v diesel as the sole carbon source, and incubated at 30°C and 250 rpm for 7 days. Cultures that developed visible and stable emulsions at the diesel-medium interphase were selected and plated onto Luria Broth (LB) agar plates. Isolated strains, showing different colony morphology, were selected, purified in axenic cultures, and stored at -80°C with the addition of glycerol 20% until use as stock culture. For biosurfactant production, each stock culture was plated in LB agar, and 2 to 5 colonies were inoculated in 10 ml LB medium and incubated overnight at 30°C and 150 rpm. This seed culture was used to inoculate 100 ml of E2 medium supplemented with diesel (10%v/v), glucose (1% w/v), or raw sunflower oil (10% v/v) as the sole carbon source, at an OD_600_ of 0.05. Cultures were incubated at 30°C and 250 rpm from 24h to 7 days.

### 2.2. 16S rRNA gene analysis

To determine the phylogeny of the isolates, DNA from each pure culture was obtained using the EasyPure Genomic DNA kit (TransGene Biotech) following the manufacturer’s instructions, and the 16S rRNA gene was sequenced (Macrogen-Korea) using 337F and 907R universal primers as described in https://www.macrogen-europe.com/service/16s-identification. To determine phylogenetic relationships of the isolated strains, the obtained sequences were compared to the Genebank nucleotide database using the BLAST platform [19]. A phylogenetic tree was made with seventeen type species of *Pseudomonas* genus, and *Escherichia/Shigella coli ATCC 11775T X80725* was selected as an outgroup. MEGA 11 software [20] was used for sequence alignment and phylogenetic tree construction utilizing the Neighbor-Joining method and a 1000-replicate bootstrap.

### 2.3. Biosurfactant production and extraction

Biosurfactant crude extract (BCE) was obtained as described by Di Martino et al [21]. Briefly, each strain was cultured as described, and cell-free supernatant was obtained by centrifugation at 10000 rpm for 5 min. The supernatant was collected, acidified to pH 2 with 6 N HCl, incubated at 4°C overnight, and then centrifuged at 10000 rpm for 10 min. The obtained pellet was resuspended in Tris-HCl 0.1 mM pH 8 buffer and extracted three times with one volume of ethyl acetate. This BCE was then concentrated with a rotavap (Büchi 461 Water Bath) at 45°C, the remnant solid was resuspended in distilled water (2 ml for each 10 ml of initial cell-free supernatant) and conserved at -20 °C for further analysis.

As per rhamnolipids standards, a cell-free supernatant from *P. aeruginosa* PAO1 was extracted as described above.

### 2.4. Biosurfactant characterization

#### 2.4.1. Emulsifying activity

The emulsification index (EI24) was determined according to Copper and Goldenberg [22]. Briefly, 5 ml of n-hexane (MERCK) was added to the same volume of cell-free supernatant, vigorously agitated by vortex for 2 min, and left to stand at room temperature for 24 h. EI24 was defined as follows:

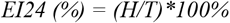

Where H is the height of the emulsion phase, and T is the total height measured in cm. As a negative control, a culture medium without inoculation was used. All measurements were run in triplicate.

#### 2.4.2. Drop collapse test

In the drop collapse test, 20 µl of cell-free supernatant was mixed with 1μl of a 0.1% (v/v) methylene blue solution. As a control, 20 µl of uncultured media was used. A 15 μl drop was placed on a hydrophobic surface (parafilm), and the contact angle was measured using ImageJ 1x software [23]. Relative contact angle (RCA) was defined as follows:

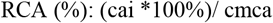

where the contact angle of the isolates (cai) was relative to the contact angle of the culture media (cmca, as control), as a total percentage (100%).

#### 2.4.3. Critical micelle concentration (CMC) of the crude extract

To obtain CMC values, the compound mass present in 10 ml of BCE was calculated as dry weight after lyophilization [24]. To determine the surface tension (ST), a Du Nouy tensiometer (Cenco Du Nouy 70545) was used. The surface tension of the distilled water used for dilutions was measured as a baseline control.

#### 2.4.4. Analysis of the chemical nature of the putative surfactants present in BCE

The chemical composition of BCE was analyzed using thin-layer chromatography (TLC) with a mobile phase consisting of chloroform, methanol, and acetic acid (65:15:2). TLCs were developed with Ninhydrin reagent to detect amino acids and peptides [25], Molisch reactive (α-naphthol and H_2_SO_4_) for detecting glycosidic moieties [26], and iodine to detect lipids moieties [27]. ImageJ 1X software was used to analyze the TLC spots [23].

### 2.5. SER assays

Soil samples were obtained from the vicinity of the industrial waste treatment plant (34° 34’ 50.7’’ S, 58° 49’ 27.2” W). 250g homogenized soil was kept in black polyethylene bags at 15°C until its subsequent use. When necessary, part of the homogenized soil was sterilized by tyndallization as described previously [28].

For microcosm construction, sterile (S) or non-sterile (NS) soil was supplemented with KNO_₃_ (2 g of N/kg of dried soil) and K_₂_HPO_₄_ (450 mg of P/kg of dried soil), adjusted to 60% field capacity, and artificially contaminated with 10% V/W diesel. Each experimental unit (ExU) consisted of the equivalent of 10 g of dried soil in a 45 mm Petri dish. Different sets of 5 ExUs each were designed using S or NS soils. When necessary, BCE was added as twice the CMC relative to the water present in the ExU (Table 1). As controls, the effect of the atenuation in diesel degradation (Table 1). To analyze whether BCE enhances nonbiological (physical-chemical) diesel degradation, sterile soil was used as a control. To analyze if the BCE enhances biological degradation, a non-sterilized soil without BCE addition was used as a control (Table 1)

**Table 1.** Description of the microcosm sets used to analyze the effect of MM and ML BCEs on diesel degradation. The table includes the tested characteristics, type of soil, and BCE added for each microcosm set.

| Microcosm set | Characteristic tested | Soil | BCE added |
| --- | --- | --- | --- |
| S1 | Abiotic degradation control | S | - |
| S2, Sn | Effect of BCE on abiotic degradation | S | + |
| NS1 | Biotic degradation control | NS | - |
| NS2, NSn | Effect of BCE on biotic degradation | NS | + |

**Table 2.**
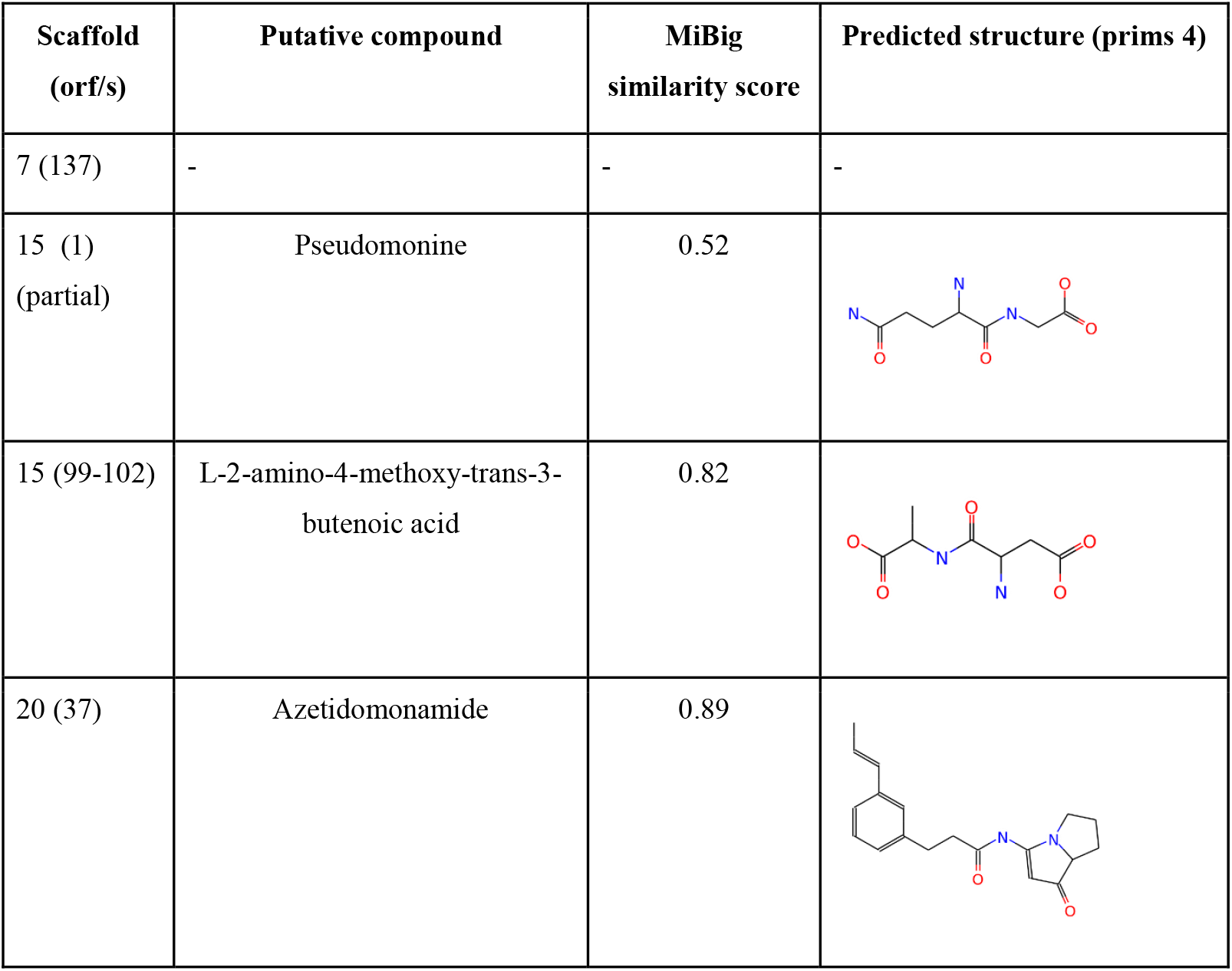
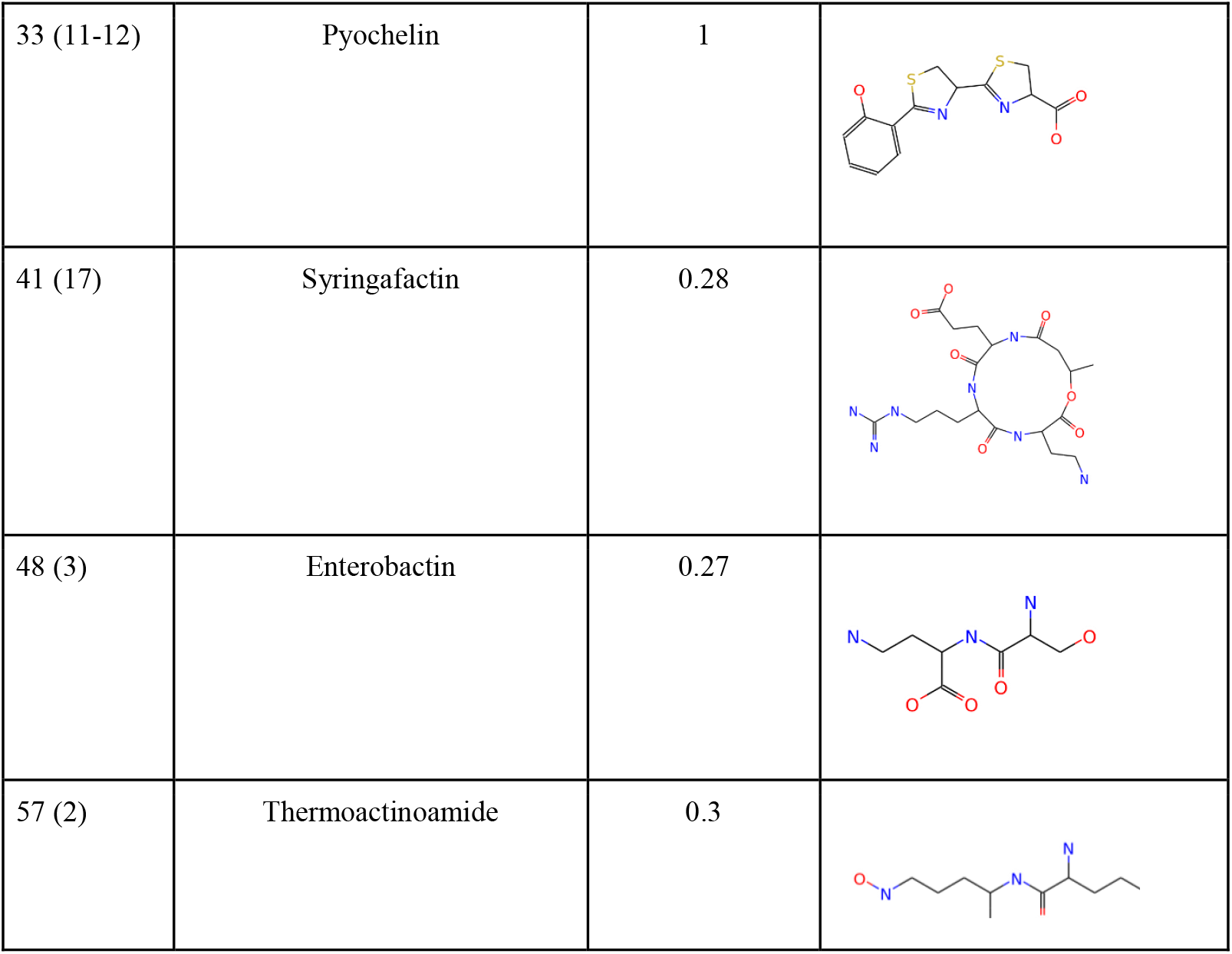
Description of scaffolds found in MM strain. The table includes the putative compound, the MiBig similarity score, and the predicted structure.

Microcosms were incubated at 24°C for 30 days with weekly agitation to enhance aerobic diesel degradation. After this time, each experimental unit was transferred into a 100 ml glass bottle, extracted with 30 ml of n-hexane: acetone 1:1, and incubated for 4 h at room temperature and 180 rpm. Two ml of the organic phase was collected and centrifuged at 12.000 g for 10 min. The clear supernatant was analyzed in a gas chromatography (Agilent 7820A GC–FID with an automatic sampler ALS 7693) using an Agilent HP-5 capillary column (30 m, 0.25 μm de weight and 0.32 mm ID) as described previously by Colonnella et al [29]. n-octane (Merck), with a retention time of approximately 6.7 min. was used as the internal control.

### 2.6. Genome annotation and bioinformatics analysis

The genome sequence was obtained by Illumina NovaSeq PE150 (Novogene Inc.). De Novo assembly was done using SOAP De Novo software, SPAdes software, and Abyss software. The results of the three software were integrated with CISA software, obtaining the 79 scaffolds. Draft genome sequence was deposited in Genebank AN: JAMWSM000000000

DNA-DNA in-silico hybridization was done using Type Strain Genome Server (https://tygs.dsmz.de) with the default setting. Biosurfactant synthesis-related genes were mined using the seed viewer platform [30], antiSMASH 7.0 bacterial version with the default settings (at https://antismash.secondarymetabolites.org), and PRISM 4 (https://prism.adapsyn.com)

### 2.7. Statistical analysis

Comparison of means between two groups was performed using the Student’s t-test. For comparing means among three or more groups, such as in the analysis of diesel degradation in microcosms under different treatments within each soil condition, one-way Analysis of Variance (ANOVA) was used. Models with and without interaction (additive) were analyzed and compared with the Akaike information criterion (AIC). The significance level used was 5%. If necessary, multiple comparisons were done with the Tukey test and Bonferroni correction. Analyses were performed using R software version 4.3.0 [31].

## 3. RESULTS

### 3.1. Characterization of the diesel-degrading, biosurfactant-producing bacteria

Five strains were isolated as described and labeled as MM, ML, B, V, and R (Supplementary Material S1). To analyze their capability to produce biosurfactants using different carbon sources, E2 medium supplemented with diesel (d), glucose (g), or sunflower oil (so) was inoculated with each strain, and the cell-free supernatant was obtained. MM and ML strains showed the lowest reduction in the relative contact angle (31, 38, and 35 for MM, and 31, 33, and 47 for ML, for diesel, sunflower oil, and glucose as carbon sources, respectively). They also showed higher EI24 values for the three tested carbon sources (32, 47, and 45 for MM, and 65, 33, and 50 for ML, for diesel, sunflower oil, and glucose as carbon sources, respectively) (Fig. 1, Supplementary Material Fig. 2.). Therefore, both strains were selected for further studies.

**Figure 1.**
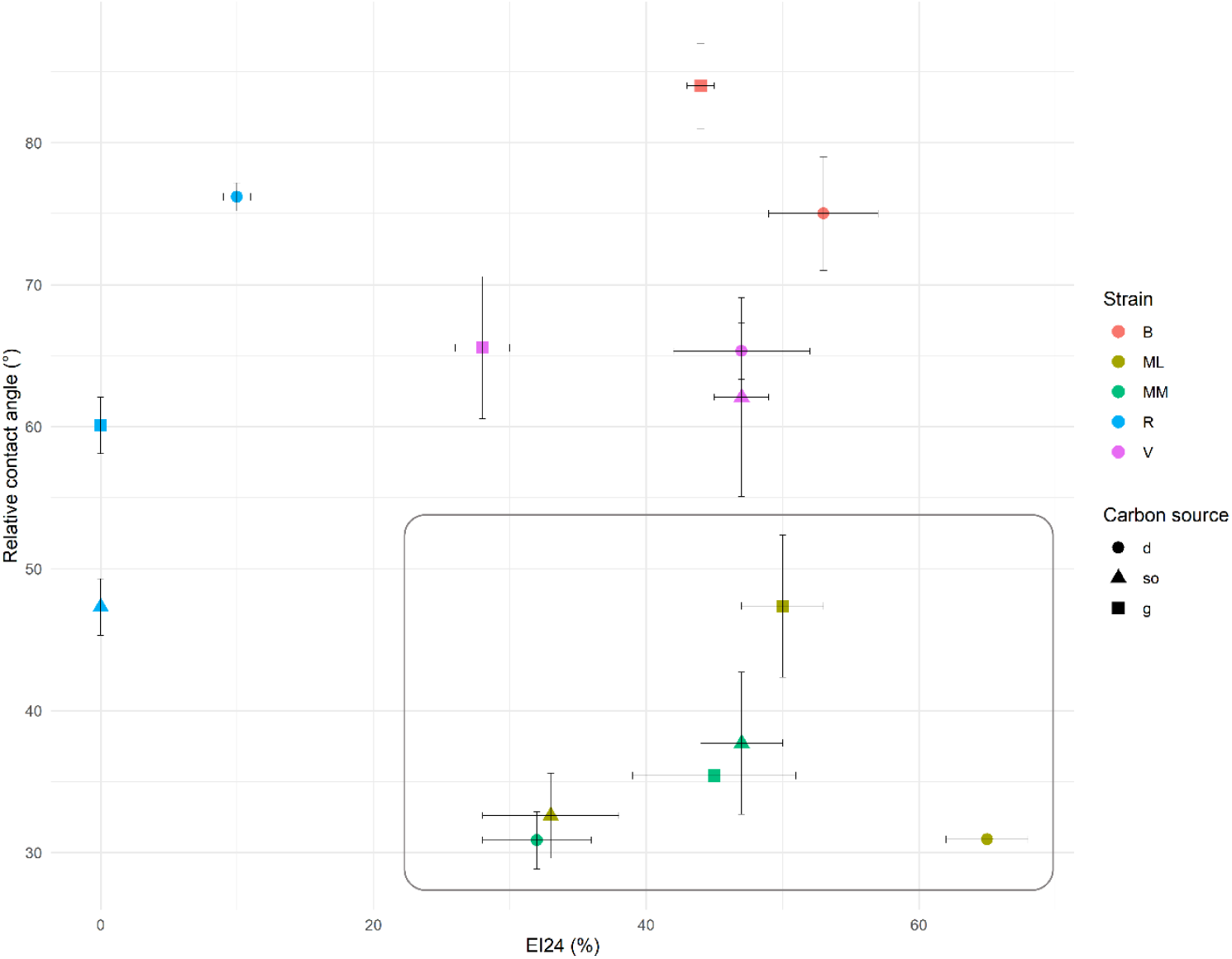
Relative contact angle values and EI24 (%) for the different isolates grown on three carbon sources (d: diesel, so: sunflower oil, g: glucose). Bars represent the standard error of three replicates. Arrows indicate the desired outcome for an effective surfactant, and the square highlights the crude extracts with the best properties.

MM and ML strains were Gram-negative, oxidase-positive, and catalase-positive. 16S rRNA gene sequencing and phylogenetic analysis showed that both are related to the *Pseudomonas aeruginosa* species (Fig. 2). BLAST comparison revealed that MM presented 99.26% identity with *Pseudomonas aeruginosa* P2 (MF289196.1) while ML showed 98.98% identity with *Pseudomonas aeruginosa* DSE2 (H1457018.1).

**Figure 2.**
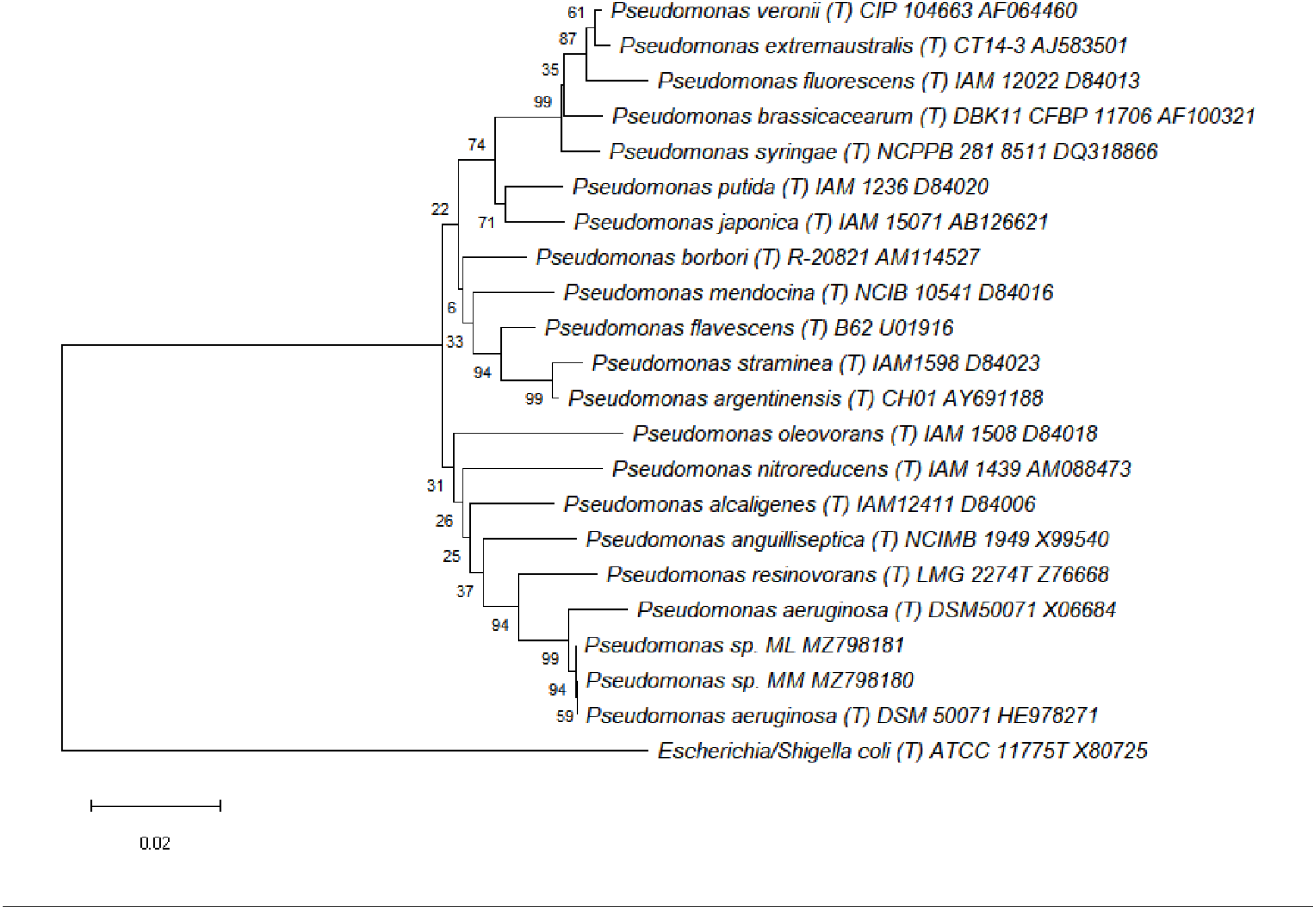
NJ phylogenetic tree consensus (Bootstrap 1000 iterations) of 16S rRNA sequences compared with type (T) *Pseudomonas* species. The sequence accession numbers are located on the right side.

### 3.2. Obtaining and Characterization of Biosurfactant Crude Extract

As described in Fig. 1 and Supplementary material Figs 1 and 2, MM and ML strains were able to produce surface-active compounds using different carbon sources. For BCE production, raw sunflower oil was chosen.

To analyze the chemical nature of the produced biosurfactant, MM-BCE and ML-BCE were assayed by thin-layer chromatography. Both BCE showed 2 spots with similar RF to *P. aeruginosa* PAO1 mono- and di-rhamnolipids, having an Rf of 0.88 and 0.47, respectively. The di/mono rhamnolipid ratio (Drh/Mrh) obtained by TLC image analysis was: MM-BCE= 2.21, ML-BCE= 0.88, PAO-BCE = 5.12.

Interestingly, MM-BCE showed a spot with aminoacidic moieties (Fig. 3A, Rf = 0.26). To determine whether the ninhydrin-stained spot corresponded to a putative lipopeptide, lipid moieties were visualized using iodine vapour, revealing a brown spot at the same Rf (Fig. 3B).

**Figure 3.**
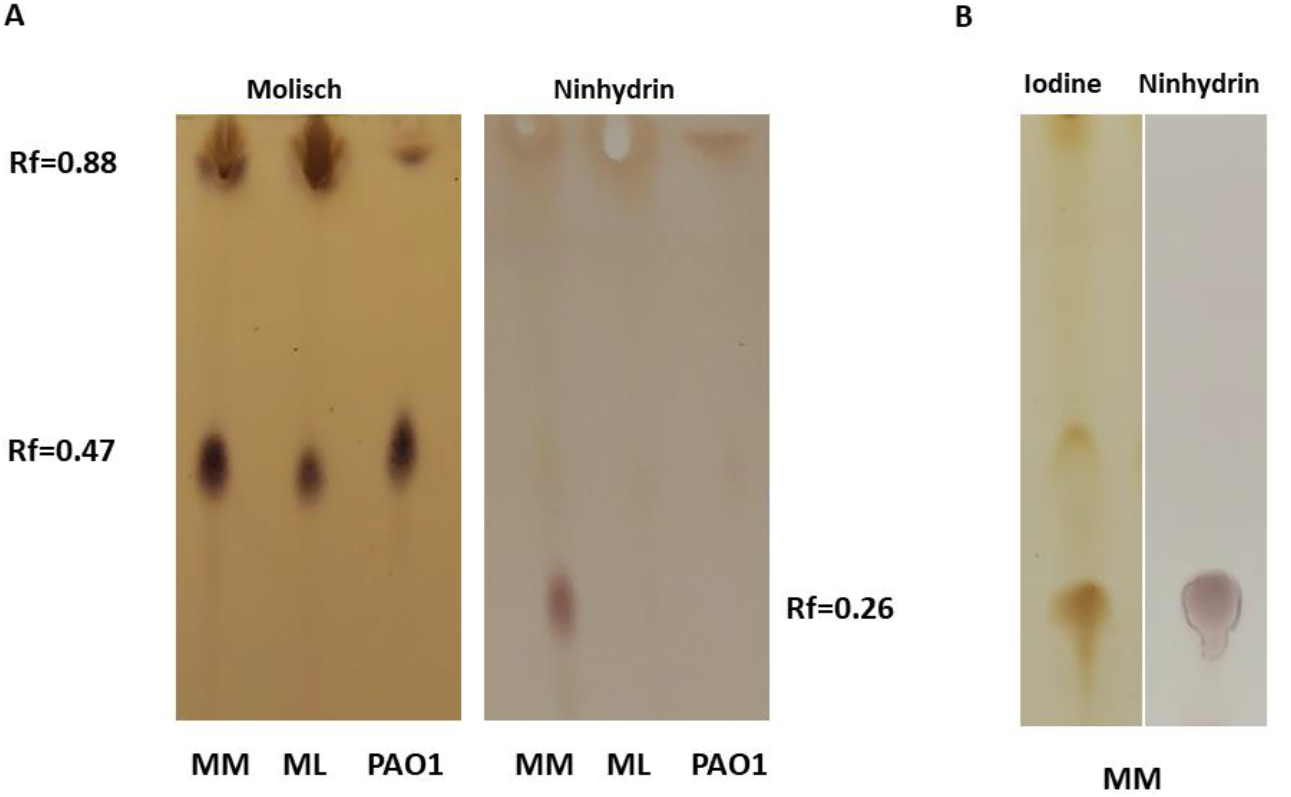
A: TLC for MM-BCE, ML-BCE, and PAO-BCE, developed with Molisch (glycolipid compounds) and Ninhydrin (peptidic compounds). B: TLC for MM-BCE developed with iodine vapour and Ninhidryn for lipids and peptides detection.

Critical micelle concentrations (CMC) of MM-BCE and ML-BCE were assayed. MM-BCE showed a CMC of 317±11ug/ml, reducing the surface tension from 72 dyn/cm to 33.5 ± 0.1, while the CMC of ML-BCE was 612±8ug/ml, producing an ST reduction from 72 dyn/cm to 33.7 ± 0.2.

### 3.3. Surfactant-enhanced remediation assay

To analyze the effect of MM-BCE and ML-BCE in diesel degradation, 6 microcosm sets were used following the setup described in Materials and Methods.

After 30 days, the remaining diesel in all sterile soil (S) microcosms was higher than in nonsterile soil (NS) microcosms (Fig. 4). No significant differences were detected among the treatments in S microcosms. When MM-BCE was added to NS, the remaining hydrocarbon decreased by 47% compared with the control (P < 0.0001, Fig. 4). On the other hand, ML-BCE showed a slight but non-significant decrease in the remaining diesel (13%). As expected, no differences were observed when the crude extract was added to S microcosms.

**Figure 4.**
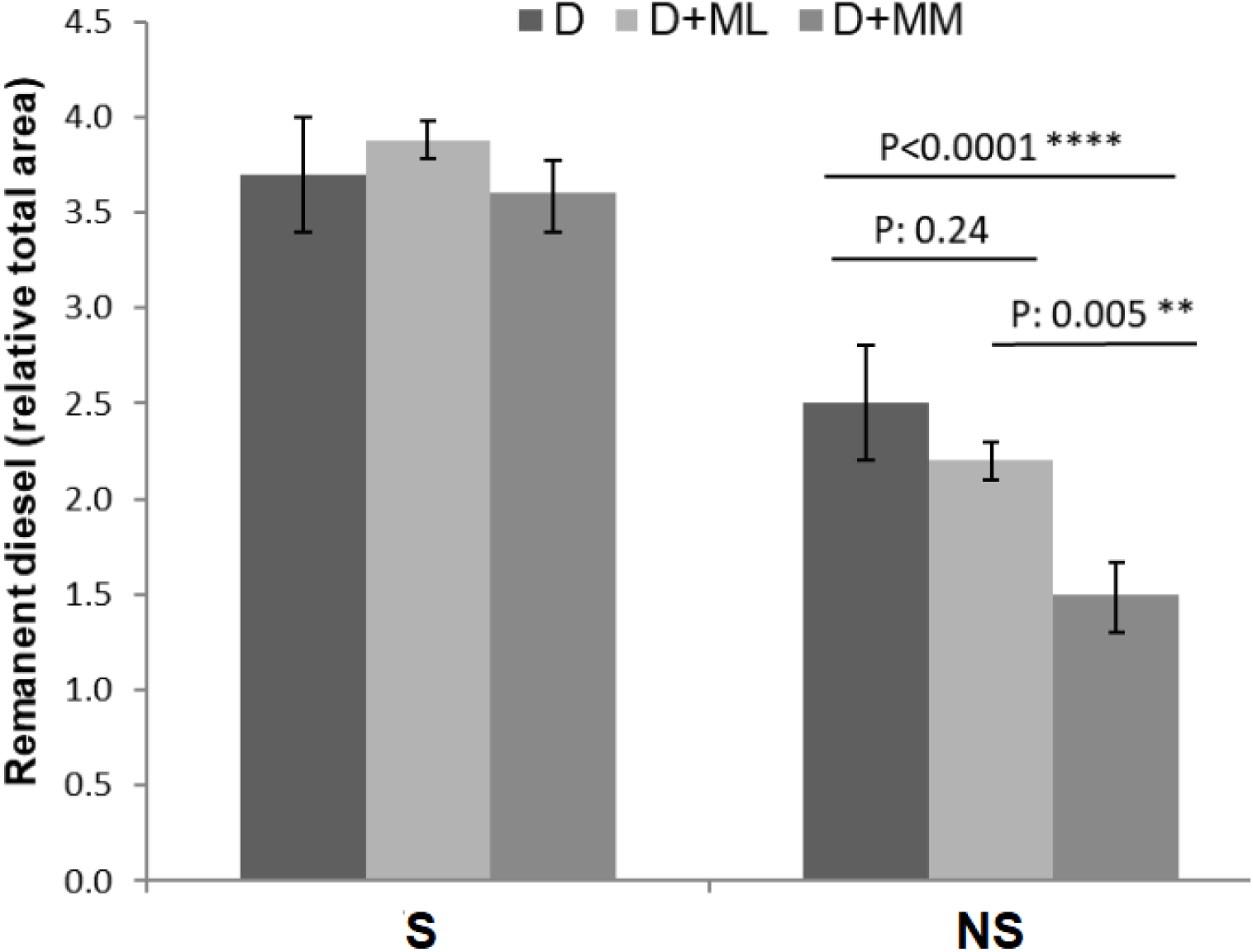
Remanent diesel (relative total area) in the S and NS microcosms with the three treatments: D for only diesel, D+MM for diesel with CEB MM, and D+ML for diesel with CEB ML.

### 3.4. Bioinformatic analysis of MM strain

Due to the MM strain showing an unexpected blend of biosurfactants with promising use in SER approaches, its draft genome sequence was obtained. It showed that the MM strain has a dDDH value of 84.2 [CI: 80.4 - 87.3] with *P. aeruginosa* DSM 50071 confirming it belongs to *P. aeruginosa* group.

The genome analysis showed the presence of the *rhl*ABRI operon in scaffold 54 with an identity of 99% with *rhl*A*BRI* operon of *P. aeruginosa* PAO1 and *rhl*C gene in scaffold 6 with an identity of 100% with *rhl*C of *P. aeruginosa* PAO1.

To search for putative NRPS that could be related to the synthesis of the lipopeptidic compound obtained in the CE, the draft genome sequence was also analyzed using Antismash and Prims 4 programs, detecting 8 putative NRPS in scaffolds 7, 15, 20, 31, 41,48, and 57 (Supplementary Material Antismash Results).

## 4. DISCUSSION

In this study, we obtained two diesel-degrading bacteria, out of 5 isolates, capable of synthesizing different biosurfactant compounds. It has been previously described that a hydrophobic carbon source increases biosurfactant production yield [32]. Because of that, we used diesel or raw sunflower oil as carbon sources for biosurfactant production. However, we finally selected sunflower oil as the carbon source to avoid the downstream purification of undesired toxic compounds from the crude biosurfactant extract.

Both selected strains belonged to the *Pseudomonas aeruginosa* species assigned by 16S rRNA gene sequencing. TLC analysis of the BCEs obtained from *P. aeruginosa* MM and ML displayed the two characteristic spots corresponding to mono- and di-rhamnolipids congeners, where the amount of di-rhamnolipids was always higher than that of mono-rhamnolipids. The difference in the RFs from the rhamnolipids spots of MM-BCE, ML-BCE, and PAO-BCE could be related to different compound congeners present in the mono- and di-rhamnolipids pool [33]. Our results also showed that MM-BCE presented a higher di-to mono-rhamnolipid ratio than ML-BCE. Although it was described that di-rhamnolipids have stronger surface activity while mono-rhamnolipids have better emulsifying activity [34], we observed that both MM-BCE and ML-BCE lowered the ST to very similar levels. Interestingly, MM-BCE also presented a spot detected by both ninhydrin and iodine staining. Although TLC cannot be use to definitively establish chemical structure, this result -combined with an extraction protocol selective for amphipathic molecules-suggests that this compound could be considered as a lipopeptide biosurfactant [35]. Lipopeptides are scarcely documented in *Pseudomonas aeruginosa* strains, and in the few described cases, species assignment was based on 16S rRNA gene sequencing [36, 37]. Here, we present for the first time a *Pseudomonas aeruginosa* strain confirmed by DNA-DNA hybridization that simultaneously produces rhamnolipids and a putative lipopeptide surfactant. The analysis of the draft genome of *P. aeruginosa* MM confirmed its capacity to produce mono- and di-rhamnolipids, as well as the presence several NRPS including a putative syringafactin synthase codified in scaffold 41. Syringafactin is one of the 4 kinds of lipopeptide surfactants described in the *Pseudomonas* genus [12]; however, to the best of our knowledge, this has not been previously described in *P. aeruginosa*. Although this synthase was identified in the genome of P. aeruginosa MM, we cannot conclusively determine whether the corresponding lipopeptide is indeed syringafactin, given the structural complexity of NRPSs, which limits the accuracy of product prediction [38]. Furthermore, we We cannot exclude the possibility that the compound we detected could be another kind of surfactant, like a lipoamino acid. Neve et al. [39] described acyl-putrescine as a rare secondary metabolite produced by very few *P. aeruginosa* isolates (less than 5%). However, lipoamino acids have been reported as membrane components, and their isolation typically requires bacterial pellets rather than cell-free supernatants [40].

To compare the surfactant performance, one of the key parameters to analyze is the critical micelle concentration (CMC) [41]. The CMC varies depending on the set of compounds produced, the carbon source, the bacterial species, and the degree of surfactant purification [32]. It was described that, in a mixture of surfactants, the CMC could be influenced by the synergism or antagonism between its compounds, with synergism being the most frequently observed effect [42]. As a result, different types of surfactants are used in many industrial products and processes instead of pure components [43, 44]. In this work, MM-BCE showed a lower CMC compared to ML-BCE. This difference could be attributed to several factors: the differences between mono- and di-rhamnolipids ratios and/or in biosurfactant composition, especially the presence of the putative lipopeptide surfactant detected. The obtained CMC values for MM-BCE and ML-BCE are consistent with the ones reported previously [45-47]. When compared with synthetic surfactants the CMC of MM-BCE was approximately sevenfold smaller than SDS [48]. Finally, the EI24 of MM-BCE and ML-BCE were similar to those previously reported [49]. Regarding surface tension reduction, both MM and ML achieved a value of 33 mN/m which is similar to what is found in the literature regarding different *P. aeruginosa* strains, ranging from 29 to 40.5 mN/m [50, 51].

The effect of the addition of biosurfactants in SER assays depends on different parameters, like the microcosms’ incubation time, the enhanced hydrocarbon bioavailability, and the effect of these compounds on the autochthonous microbiota.

Regarding the incubation time, higher percentages (80-85%) were observed after 6 months [52] while Akbari et al [53] reported about 10% of total hydrocarbon reduction after 30 incubation days and 20% after 80 days when rhamnolipids were added to microcosm assays. This suggests that the MM-BCE is effective and competitive, having a degradation rate of 47% after 30 days.

To distinguish between enhanced hydrocarbon (HC) bioavailability and the changes in the microbial community induced by biosurfactants, we included both sterile (abiotic) and non-sterile (biotic) soils as controls. Our results, by comparing sterile soil with or without BCEs, showed that surfactant addition does not affect diesel attenuation. But, when the native microbial communities were present, our results showed that MM-BCE and ML-BCE addition increased the rate of hydrocarbon degradation, but with different effectiveness. It was previously described that different mixtures of surface-active compounds may have different capabilities to enhance the remediation of hydrophobic pollutants [54]. Szulc et al [52] found no significant effect when commercial rhamnolipids were used, while Cameotra and Singh [55] had a degradation of 95% using crude biosurfactants by *Pseudomonas aeruginosa* and *Rhodococcus* sp. In this work, the presence of a blend of rhamnolipids and lipopeptides in BCE-MM could be responsible for its best performance. The use of surfactants with diverse characteristics has demonstrated positive effects in previous works. For instance, Cheng et al. [56] showed that a blend of glycolipid and lipopeptide produced by *B. subtilis* strain achieved 86% oil-washing efficiency. Their findings revealed that the lipopeptide primarily contributes to stability and oil-washing efficiency, while the glycolipid plays a greater role in enhancing emulsifying activity.

The differences observed between the addition of MM-BCE and ML-BCE could be due not only to an increase in hydrocarbon bioavailability but also to how each BCE affected the bacterial community, especially the hydrocarbon-degrading taxa [57, 58]. In line with the study by Feng et al. [59], biosurfactants such as sophorolipids can significantly improve the bioavailability of total petroleum hydrocarbons (TPH) by enhancing their desorption from soil particles and facilitating transmembrane transport. Additionally, biosurfactants can act as co-substrates, supporting microbial growth and co-metabolic degradation of hydrocarbons. These compounds were also shown to increase cell surface hydrophobicity and stimulate key enzymatic activities and the expression of functional genes related to TPH degradation.

## 5. CONCLUSION

In this work, we present for the first time a fully sequenced *P. aeruginosa* strain capable of simultaneously synthesizing a blend of glycolipids and a putative lipopeptides. A crude extract of this surfactant mix, with minimal post-production purification, resulted in significant diesel biodegradation compared to the controls in microcosm assays. These results represent a promising amendment to be used in the bioremediation of petroleum-contaminated sites and microbial-enhanced oil recovery (MEOR).

## FUNDING AND COMPETING INTERESTS

This work was supported by a grant from the National Scientific and Technical Research Council - Argentina (Argentina) (Grant N°. PUE14120210300619CO, PIP 11220200101436CO)

LJRI is a career investigator from CONICET and the University of Buenos Aires

The authors have no competing interests to declare that are relevant to the content of this article

## ACKNOWLEDGMENTS

The author wants to thank Hugo Sato for his assistance with the illustrations, and Ruby Terranova and Marcos Dennin for their technical assistance.

## REFERENCES

1. Sakhaei, Z., Riazi, M., 2022. In-situ petroleum hydrocarbons contaminated soils remediation by polymer enhanced surfactant flushing: Mechanistic investigation. Process Saf. Environ. Protect. 161, 758–770. 10.1016/j.psep.2022.03.086

2. Kebede, G., Tafese, T., Abda, E.M., Kamaraj, M., Assefa, F., 2021. Factors influencing the bacterial bioremediation of hydrocarbon contaminants in the soil: mechanisms and impacts. J. Chem. 2021(1), 9823362. 10.1155/2021/9823362

3. IARC Working Group on the Evaluation of Carcinogenic Risks to Humans, 1989. Diesel fuels. In: Occupational Exposures in Petroleum Refining; Crude Oil and Major Petroleum Fuels. International Agency for Research on Cancer (IARC). https://www.ncbi.nlm.nih.gov/books/NBK531266/

4. Mohanta, S., Pradhan, B., Behera, I.D., 2024. Impact and remediation of petroleum hydrocarbon pollutants on agricultural land: a review. Geomicrobiol. J. 41(4), 345–359. 10.1080/01490451.2023.2243925

5. Sui, X., Wang, X., Li, Y., Ji, H., 2021. Remediation of petroleum-contaminated soils with microbial and microbial combined methods: Advances, mechanisms, and challenges. Sustainability 13(16), 9267. 10.3390/su13169267

6. Mohanty, S., Jasmine, J., Mukherji, S., 2013. Practical considerations and challenges involved in surfactant enhanced bioremediation of oil. BioMed Res. Int. 2013(1), 328608. 10.1155/2013/328608

7. Mulligan, C.N., Yong, R.N., Gibbs, B.F., 2001. Surfactant-enhanced remediation of contaminated soil: a review. Eng. Geol. 60(1–4), 371–380. 10.1016/S0013-7952(00)00117-4

8. Johnson, P., Trybala, A., Starov, V., Pinfield, V.J., 2021. Effect of synthetic surfactants on the environment and the potential for substitution by biosurfactants. Adv. Colloid Interface Sci. 288, 102340. 10.1016/j.cis.2020.102340

9. Kashif, A., Rehman, R., Fuwad, A., Shahid, M.K., Dayarathne, H.N.P., Jamal, A., …, Choi, Y., 2022. Current advances in the classification, production, properties and applications of microbial biosurfactants—A critical review. Adv. Colloid Interface Sci. 306, 102718. 10.1016/j.cis.2022.102718

10. Thavasi, R., Banat, I.M., 2019. Introduction to microbial biosurfactants. In: Microbial Biosurfactants and their Environmental and Industrial Applications. CRC Press, 1–15. 10.1201/b21950

11. Hayes, D.G., Solaiman, D.K., Ashby, R.D., 2019. Biobased Surfactants: Synthesis, Properties, and Applications. Elsevier.

12. Chauhan, V., Mazumdar, S., Pandey, A., Kanwar, S.S., 2023. Pseudomonas lipopeptide: An excellent biomedical agent. MedComm–Biomater. Appl. 2(1), e27. 10.1002/mba2.27

13. Sarubbo, L.A., da Gloria, C.S., Durval, I.J.B., Bezerra, K.G.O., Ribeiro, B.G., Silva, I.A., …, Banat, I.M., 2022. Biosurfactants: Production, properties, applications, trends, and general perspectives. Biochem. Eng. J. 181, 108377. 10.1016/j.bej.2022.108377

14. Zhuang, X., Wang, Y., Wang, H., Dong, Y., Li, X., Wang, S., …, Wu, S., 2022. Comparison of the efficiency and microbial mechanisms of chemical- and bio-surfactants in remediation of petroleum hydrocarbon. Environ. Pollut. 314, 120198. 10.1016/j.envpol.2022.120198

15. Eras-Muñoz, E., Farré, A., Sánchez, A., Font, X., Gea, T., 2022. Microbial biosurfactants: a review of recent environmental applications. Bioengineered 13(5), 12365–12391. 10.1080/21655979.2022.2074621

16. Invally, K., Sancheti, A., Ju, L.K., 2019. A new approach for downstream purification of rhamnolipid biosurfactants. Food Bioprod. Process. 114, 122–131. 10.1016/j.fbp.2018.12.003

17. Jiang, J., Zu, Y., Li, X., Meng, Q., Long, X., 2020. Recent progress towards industrial rhamnolipids fermentation: process optimization and foam control. Bioresour. Technol. 298, 122394. 10.1016/j.biortech.2019.122394

18. Lageveen, R.G., Huisman, G.W., Preusting, H., Ketelaar, P., Eggink, G., Witholt, B., 1988. Formation of polyesters by Pseudomonas oleovorans: effect of substrates on formation and composition of poly-(R)-3-hydroxyalkanoates and poly-(R)-3-hydroxyalkenoates. Appl. Environ. Microbiol. 54(12), 2924–2932. 10.1128/aem.54.12.2924-2932.1988

19. Altschul, S.F., Gish, W., Miller, W., Myers, E.W., Lipman, D.J., 1990. Basic local alignment search tool. J. Mol. Biol. 215(3), 403–410. 10.1016/S0022-2836(05)80360-2

20. Tamura, K., Stecher, G., Kumar, S., 2021. MEGA11: Molecular evolutionary genetics analysis version 11. Mol. Biol. Evol. 38(7), 3022–3027. 10.1093/molbev/msab120

21. Di Martino, C., López, N.I., Raiger Iustman, L.J., 2012. Isolation and characterization of benzene, toluene and xylene degrading Pseudomonas sp. selected as candidates for bioremediation. Int. Biodeterior. Biodegrad. 67, 15–20. 10.1016/j.ibiod.2011.11.004

22. Cooper, D.G., Goldenberg, B.G., 1987. Surface-active agents from two Bacillus species. Appl. Environ. Microbiol. 53(2), 224–229. 10.1128/aem.53.2.224-229.1987

23. Schneider, C.A., Rasband, W.S., Eliceiri, K.W., 2012. NIH Image to ImageJ: 25 years of image analysis. Nat. Methods 9(7), 671–675. 10.1038/nmeth.2089

24. Duvnjak, Z., Cooper, D.G., Kosaric, N., 1982. Production of surfactant by Arthrobacter paraffineus ATCC 19558. Biotechnol. Bioeng. 24(1), 165–175. 10.1002/bit.260240114

25. Baranowska, I., Kozłowska, M., 1995. TLC separation and derivative spectrophotometry of some amino acids. Talanta 42(10), 1553–1557. 10.1016/0039-9140(95)01569-W

26. Taki, T., Ishikawa, H., Imai, K., Yachi, A., Matsumoto, M., 1985. Immunological analysis of glycolipids on cell surfaces of cultured human tumor cell lines: expression of lactoneotetraosylceramide on tumor cell surfaces. J. Biochem. 98(4), 887–895. 10.1093/oxfordjournals.jbchem.a135368

27. Barale, S.S., Ghane, S.G., Sonawane, K.D., 2022. Purification and characterization of antibacterial surfactin isoforms produced by Bacillus velezensis SK. AMB Express 12(1), 7. 10.1186/s13568-022-01348-3

28. MacRae, I.C., Raghu, K., Castro, T.F., 1967. Persistence and biodegradation of four common isomers of benzene hexachloride in submerged soils. J. Agric. Food Chem. 15(5), 911–914. 10.1021/jf60153a030

29. Colonnella, M.A., Lizarraga, L., Rossi, L., Díaz Peña, R., Egoburo, D., López, N.I., Raiger Iustman, L.J., 2019. Effect of copper on diesel degradation in Pseudomonas extremaustralis. Extremophiles 23, 91–99. 10.1007/s00792-018-1063-2

30. Overbeek, R., 2005. The subsystems approach to genome annotation and its use in the project to annotate 1000 genomes. Nucleic Acids Res. 33(17), 5691–5702. 10.1093/nar/gki866

31. R Development Core Team, 2024. R: A Language and Environment for Statistical Computing. R Foundation for Statistical Computing, Vienna, Austria. https://www.R-project.org

32. de Oliveira Schmidt, V.K., de Souza Carvalho, J., de Oliveira, D., Andrade, C.J., 2021. Biosurfactant inducers for enhanced production of surfactin and rhamnolipids: an overview. World J. Microbiol. Biotechnol. 37, 21. 10.1007/s11274-020-02970-8

33. El-Housseiny, G.S., Aboshanab, K.M., Aboulwafa, M.M., Hassouna, N.A., 2020. Structural and physicochemical characterization of rhamnolipids produced by Pseudomonas aeruginosa P6. AMB Express 10(1), 201. 10.1186/s13568-020-01141-0

34. Zhao, F., Shi, R., Ma, F., Han, S., Zhang, Y., 2018. Oxygen effects on rhamnolipids production by Pseudomonas aeruginosa. Microb. Cell Fact. 17, 1–11. 10.1186/s12934-018-0888-9

35. Dini, S., Oz, F., Bekhit, A. E. D. A., Carne, A., & Agyei, D. (2024). Production, characterization, and potential applications of lipopeptides in food systems: A comprehensive review. Compr. Rev. Food Sci. Food Saf. 23(4), e13394. 10.1111/1541-4337.13394

36. Liu, W.J., Duan, X.D., Wu, L.P., Masakorala, K., 2018. Biosurfactant production by Pseudomonas aeruginosa SNP0614 and its effect on biodegradation of petroleum. Appl. Biochem. Microbiol. 54, 155–162. 10.1134/S0003683818020060

37. Bezza, F.A., Chirwa, E.M.N., 2016. Biosurfactant-enhanced bioremediation of aged polycyclic aromatic hydrocarbons (PAHs) in creosote contaminated soil. Chemosphere 144, 635–644. 10.1016/j.chemosphere.2015.08.027

38. Duban, M., Cociancich, S., & Leclère, V. (2022). Nonribosomal peptide synthesis definitely working out of the rules. Microorganisms, 10(3), 577.

39. Neve, R.L., Giedraitis, E., Akbari, M.S., Cohen, S., Phelan, V.V., 2024. Secondary metabolite profiling of Pseudomonas aeruginosa isolates reveals rare genomic traits. mSystems 9(5), e00339–24. 10.1128/msystems.00339-24

40. Hueber, A., Petitfils, C., Le Faouder, P., Langevin, G., Guy, A., Galano, J.M., …, Bertrand-Michel, J., 2022. Discovery and quantification of lipoamino acids in bacteria. Anal. Chim. Acta 1193, 339316. 10.1016/j.aca.2021.339316

41. Kim, J., Vipulanandan, C., 2006. Removal of lead from contaminated water and clay soil using a biosurfactant. J. Environ. Eng. 132(7), 777–786. 10.1061/(ASCE)0733-9372(2006)132:7(777)

42. Rosen, M.J., Kunjappu, J.T., 2012. Surfactants and Interfacial Phenomena (4th ed.). Wiley. 10.1002/9781118228920

43. Shah, M.U.H., Reddy, A.V.B., Yusup, S., Goto, M., Moniruzzaman, M., 2021. Ionic liquid-biosurfactant blends as effective dispersants for oil spills: Effect of carbon chain length and degree of saturation. Environ. Pollut. 284, 117119. 10.1016/j.envpol.2021.117119

44. Singh, R.D., Ganesan, N.G., Rangarajan, V., 2023. A biosurfactant cocktail-based formula for the formulation of stable skin-care cosmetic nanoemulsion. Colloid J. 85(3), 442–455. 10.1134/S1061933X23600082

45. Bordas, F., Lafrance, P., Villemur, R., 2005. Conditions for effective removal of pyrene from an artificially contaminated soil using Pseudomonas aeruginosa 57SJ rhamnolipids. Environ. Pollut. 138(1), 69–76. 10.1016/j.envpol.2005.02.017

46. Pornsunthorntawee, O., Wongpanit, P., Chavadej, S., Abe, M., Rujiravanit, R., 2008. Structural and physicochemical characterization of crude biosurfactant produced by Pseudomonas aeruginosa SP4 isolated from petroleum-contaminated soil. Bioresour. Technol. 99(6), 1589–1595. 10.1016/j.biortech.2007.04.020

47. Silva, S.N.R.L., Farias, C.B.B., Rufino, R.D., Luna, J.M., Sarubbo, L.A., 2010. Glycerol as substrate for the production of biosurfactant by Pseudomonas aeruginosa UCP0992. Colloids Surf. B Biointerfaces 79(1), 174–183. 10.1016/j.colsurfb.2010.03.050

48. Valsaraj, K.T., Gupta, A., Thibodeaux, L.J., Harrison, D.P., 1988. Partitioning of chloromethanes between aqueous and surfactant micellar phases. Water Res. 22(9), 1173–1183. 10.1016/0043-1354(88)90013-9

49. Pourfadakari, S., Ghafari, S., Takdastan, A., Jorfi, S., 2021. A salt resistant biosurfactant produced by moderately halotolerant Pseudomonas aeruginosa (AHV-KH10) and its application for bioremediation of diesel-contaminated sediment in saline environment. Biodegradation 32, 327–341. 10.1007/s10532-021-09941-2

50. Rikalovic, M.G., Abdel-Mawgoud, A.M., Déziel, E., Gojgic-Cvijovic, G.D., Nestorovic, Z., Vrvic, M.M., Karadzic, I.M., 2013. Comparative analysis of rhamnolipids from novel environmental isolates of Pseudomonas aeruginosa. J. Surfactants Deterg. 16(5), 673–682. 10.1007/s11743-013-1462-4

51. Costa, S.G., Nitschke, M., Lépine, F., Déziel, E., Contiero, J., 2010. Structure, properties and applications of rhamnolipids produced by Pseudomonas aeruginosa L2-1 from cassava wastewater. Process Biochem. 45(9), 1511–1516. 10.1016/j.procbio.2010.05.033

52. Szulc, A., Ambrożewicz, D., Sydow, M., Ławniczak, Ł., Piotrowska-Cyplik, A., Marecik, R., Chrzanowski, Ł., 2014. The influence of bioaugmentation and biosurfactant addition on bioremediation efficiency of diesel-oil contaminated soil: feasibility during field studies. J. Environ. Manage. 132, 121–128. 10.1016/j.jenvman.2013.11.006

53. Akbari, A., Kasprzyk, A., Galvez, R., Ghoshal, S., 2021. A rhamnolipid biosurfactant increased bacterial population size but hindered hydrocarbon biodegradation in weathered contaminated soils. Sci. Total Environ. 778, 145441. 10.1016/j.scitotenv.2021.145441

54. Mendes, A.N., Filgueiras, L.A., Pinto, J.C., Nele, M., 2015. Physicochemical properties of rhamnolipid biosurfactant from Pseudomonas aeruginosa PA1 to applications in microemulsions. J. Biomater. Nanobiotechnol. 6(1), 64–79. 10.4236/jbnb.2015.61007

55. Cameotra, S.S., Singh, P., 2008. Bioremediation of oil sludge using crude biosurfactants. Int. Biodeterior. Biodegrad. 62(3), 274–280. 10.1016/j.ibiod.2007.11.009

56. Cheng, F., Tang, C., Yang, H., Yu, H., Chen, Y., Shen, Z., 2013. Characterization of a blend-biosurfactant of glycolipid and lipopeptide produced by Bacillus subtilis TU2 isolated from underground oil-extraction wastewater. J. Microbiol. Biotechnol. 23(3), 390–396. 10.4014/jmb.1207.09020

57. Lu, L., Zhang, J., Peng, C., 2019. Shift of soil polycyclic aromatic hydrocarbons (PAHs) dissipation pattern and microbial community composition due to rhamnolipid supplementation. Water Air Soil Pollut. 230, 1–14. 10.1007/s11270-019-4118-9

58. Posada-Baquero, R., Jiménez-Volkerink, S.N., García, J.L., Vila, J., Cantos, M., Grifoll, M., Ortega-Calvo, J.J., 2020. Rhizosphere-enhanced biosurfactant action on slowly desorbing PAHs in contaminated soil. Sci. Total Environ. 720, 137608. 10.1016/j.scitotenv.2020.137608

59. Feng, L., Jiang, X., Huang, Y., Wen, D., Fu, T., Fu, R., 2021. Petroleum hydrocarbon-contaminated soil bioremediation assisted by isolated bacterial consortium and sophorolipid. Environ. Pollut. 273, 116476. 10.1016/j.envpol.2021.116476

